# Elucidating the structural features of ABCA1 in its heterogeneous membrane environment

**DOI:** 10.1101/2021.05.23.445323

**Authors:** Sunidhi S, Sukriti Sacher, Parth Garg, Arjun Ray

## Abstract

ATP Binding Cassette Transporter A1 (ABCA1) plays an integral part in Reverse Cholesterol Transport (RCT) and is critical for maintaining lipid homeostasis. One theory of lipid efflux by the transporter (alternating access) proposes that ABCA1 harbours two different conformations that provide alternating access for lipid binding and release. This is followed by a sequestration via a direct interaction between ABCA1 and its partner, ApoA1. The alternative theory (lateral access) proposes that ABCA1 obtains lipids laterally from the membrane to form a temporary extracellular “reservoir”. This reservoir contains an isolated lipid monolayer due to the net accumulation of lipids in the exofacial leaflet. Recently, a full-length Cryo-EM structure of this 2,261-residue transmembrane protein showed its discreetly folded domains and have detected the presence of a tunnel enclosed within ECDs but not in theTMDs giving it an outward-facing conformation. This structure was hypothesized to substantiate the lateral access theory, whereby ApoA1 obtained lipids from the proximal end laterally. Utilizing long time-scale multiple replica atomistic molecular dynamics simulations (MDS), we simulated the structure in a large heterogeneous lipid environment and found that along with several large conformational changes, the protein harbours a continuous tunnel that traverses the entire length of the protein. In this study, we have characterized ABCA1 and the lipid dynamics along with the protein-lipid interactions in the heterogeneous environment, providing novel insights into understanding ABCA1 conformation at an atomistic level.

---

ATP Binding Cassette transporters constitute a ubiquitous superfamily of integral membrane proteins that couple transport of a chemically diverse set of substrates across bilayers to hydrolysis of ATP. The ABCA subfamily that includes ABCA1, is involved in transportation of lipids across the membrane(1). ABCA1 plays a pivotal role in Reverse Cholesterol Transport (RCT) by facilitating the unidirectional net efflux of free cholesterol and phospholipids to poorly lipidated ApoA1 in plasma. This affiliation leads to the formation of nascent HDL(2). The physiological importance of ABCA1 is underscored by its association with various dyslipidaemic disorders such as familial HDL deficiency and Tangier disease, both of which are characterized by accumulation of cholesteryl esters in peripheral cells, particularly macrophages(3). Moreover, ABCA1-mutant mice show diminished LCAT activity, lack of α-migrating HDL particles and triglyceride-rich pre-β HDL particles, highlighting its role in nascent HDL formation(4, 5). This is also supported by *in-vitro* models that have concluded a consequent failure in nascent HDL generation on incubation of ApoA1 with cells lacking in ABCA1(2).

ABCA1 is a full transporter made up of 2,261 residues that form six domains: two transmembrane domains (TMDs, consisting of six membrane-spanning helices each); two nucleotide binding domains (NBDs) in the cytoplasm that serve to couple ATP hydrolysis to translocase activity; and two large extracellular domains (ECDs) that are implicated in protein-protein interactions as well as regulatory roles (6) **(Fig 1A)**.

**Figure 1:**
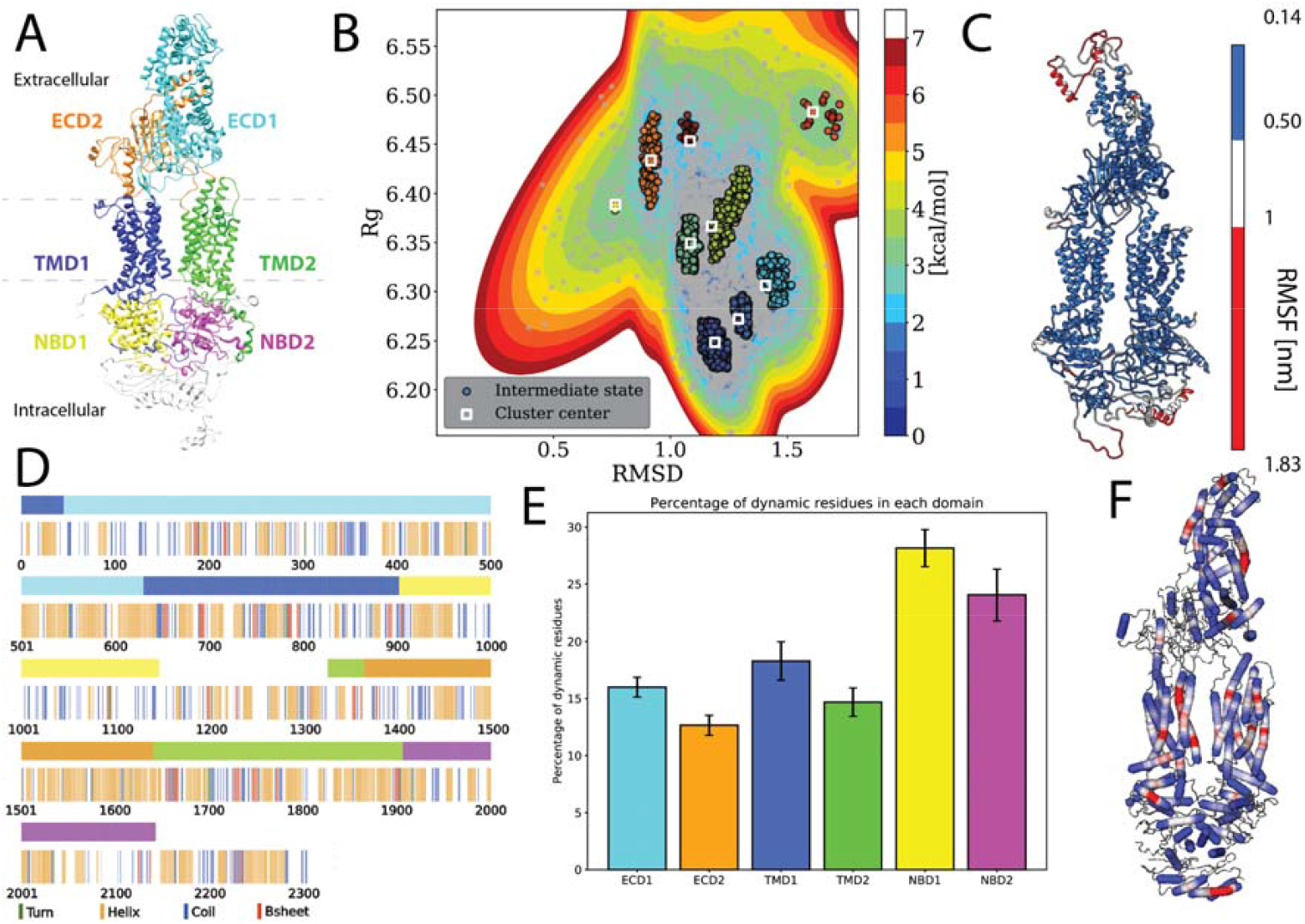
Protein specific analysis. **A**. Domain assignment of ABCA1 according to Qian et al. **B**. Free energy estimation and clustering showing distinct minima bins **C**. Root mean square fluctuation (RMSF) mapped onto the protein that showed the extremities of the protein being susceptible to large fluctuations. **D**. Residue-wise and domain-wise assignment of favoured secondary structure amongst turn (green), helix (orange), coil (blue), b-strands (red) and dynamic residues (white) according to the 1A colour scheme (top bar) **E**. Percentage of dynamic residues in each domain, where the color of each bar corresponds to the domains in Panel A. **F**. Degree of flexibility of helices constituting the protein where regions in red indicate areas of highest helix axis angle (>15□) and flexibility, white for intermediate angle magnitudes (0-15□) and blue for angles close to, or at 0□.

The two extended ECDs adopt a novel fold. ECD1 folds into 21 α helices and 10 β strands that make up 4 distinct regions: a base domain, helical domain 1 and 2 and a lid domain while ECD2 folds into 6 α helices and 7 β strands(6). The two ECDs pack tightly against each other with the help of 4 disulfide bonds (Cys75-Cys309, Cys54-Cys81, Cys355-Cys504 in ECD1 and Cys1465-Cys1477 in ECD2), two of which have been shown to be necessary for ApoA1 binding and HDL formation(7). They also enclose a predominantly hydrophobic tunnel ∼ 60 Å in height, that is open to extracellular milieu on both its distal and proximal ends. This hollow interior is conjectured to serve as a temporary storage or delivery passage for lipids (6, 8). This structure showed that the distal end of ECD1, close to the lid, forms a narrow opening that precludes passage of lipids. The proximal end however was found to be wide enough for lipids to pass. This region was therefore proposed to be the site of delivery of lipids to ApoA1, which is postulated to associate with the transmembrane protein laterally. Additionally, the proximal tunnel opening is ∼30 Å away from the upper end of the transmembrane cavity. Therefore, a pronounced conformational change (in response to ATP hydrolysis) is predicted to occur in order to facilitate the delivery of lipids from the membrane to ECDs (6, 9).

High resolution structures of NBDs of bacterial vitamin B12 importer, BtuCD as well as dimeric multidrug exporter, Sav1886 have aided in elucidation of mechanism of substrate translocation. The two-step mechanism involves binding of the two ATP molecules in the conserved Walker A and Walker B motif of NBDs. This leads to a conformational change such that the TMDs form an outward facing cavity that facilitate substrate binding. Hydrolysis of ATP puts an end to this cycle by separating the NBDs, driving TMDs to form an inward facing cavity that forms a passage for export of substrate(1). The recent Cryo EM crystal structure of a nucleotide-free ABCA1 however unveiled a snapshot of the protein such that these TMDs were found to face outwards (with a opening that faces the extracellular milieu), making ABCA1 an outlier amongst proteins of the ABC family whose structures have been captured. Moreover, this structure did not observe a continuous central cavity in the TMD region. Additionally, this ABCA1 structure also revealed a polar cluster on one side of TMD1 that was speculated to bind polar groups of lipids and facilitate their flopping movement across the hydrophobic barrier.

Although the role of ABCA1 in RCT has long been established, its functional characterization on the membrane is still in its elementary stages. It is reported that ABCA1 translocase activity leads to lipids reorganization such that free cholesterol in the membrane is more accessible to cholesterol oxidase. This reorganization is believed to be due to the partitioning of ABCA1 into non-raft domains caused by destabilization of rafts(10, 11). In addition to ApoA1 dependent export of excess cholesterol, ABCA1 is also involved in flopping cholesterol from inner to outer leaflet of the plasma membrane(12). While several studies have biochemically characterized ABCA1 activity on the membrane and its involvement in nascent HDL generation, no work has been done to explicate ABCA1’s interaction with adjacent membrane lipids at an atomistic resolution.

Transporters form a bridge between the extracellular and intracellular milieu by functioning at the junction of cell membranes. Their structural analysis however, requires them to be extracted and isolated as separate particles destroying their natural habitat(13). Several biochemical and biophysical studies have demonstrated specific protein-lipid interactions that influence assembly, stability and function of membrane proteins which remain uncharacterized due to this experimental setup (14–16). Since, the membrane imparts dramatic changes in the physical properties of the protein, it is important that they be included in models both experimental or theoretical, aimed at understanding the functional dynamics of membrane proteins. Computational methods like Molecular dynamics (MD) simulations offer a powerful complementary approach with a sufficiently high temporal and spatial resolution that allow for elucidation of atomic-level structural information and energetics of lipid-protein interactions.

In this study we have employed the use of all-atom molecular dynamics to simulate the ATP unbound ABCA1 monomer in an explicit heterogeneous membrane bilayer to refine the original model. Additionally, we have tried to assess the evidence ascertained in the Cryo-EM structure and its proposed hypothesis. We highlight the effect of immersing ABCA1 in its natural habitat on its overall architecture and volumetry as well as its secondary structural makeup. We have also quantitated the protein-lipid interactions in order to elucidate the effect of presence of protein on membrane dynamics. This study sheds light on the protein-lipid interactions that impact ABCA1 stability in the membrane.

## Results & discussion

### Stable conformation of ABCA1 in a heterogenous bilayer

Using multiple long scale (five replicates of 1μs each) all-atom classical molecular dynamics, we established the stable conformation of the protein in a heterogenous lipid membrane. The deviation from initial Cryo-EM structure is shown in **Fig S1A and S1B** that depicts the mean RMSD and Rg values of the five replicate simulations. It was observed that the final structure from the simulated trajectory stabilized with ∼1.2nm deviation on average from the initial structure. This stable structure showed an increase of 0.17 nm in its Rg value. Next, we clustered the various protein conformations that existed throughout our simulations based on the structural deviations that it underwent from its starting CryoEM state. Structural deviations characterized by the RMSD and Rg values yielded seven distinct clusters **(Fig 1B)**.

Additionally, the analysis of the mean structural fluctuations of protein throughout the simulation length revealed higher fluctuations (>1nm) in the extremities, notably in the ECD1, NBD1 and NBD2 **(Fig 1C and S2)** while the major portion of the protein remained relatively stable throughout the simulation. The domain concerted motion and PCA analysis of the trajectories also depict maximum movement in these regions **(Fig S3)**. This result is expected since both these regions are water-exposed and are known to undergo conformational changes during the ATP hydrolysis cycle(9, 17).

### Secondary structural characteristics of ATP unbound ABCA1 monomer in lipid bilayer

We utilized all the five replicates to elucidate the secondary structure assignment for each residue. In order to do this, the probability of every residue to conform to a particular secondary structural element was computed **(Fig S4)**. Based on a residue’s propensity to exist as either a helix, turn, strand or coil, for 70% of the time throughout the simulation, a secondary structure was assigned to it (**Fig S5**). It was observed that the protein structure was predominantly constituted by helices (41% residues), followed by coil (21% residues), while turns and β-strands were least favoured (14.34% and 5.41% respectively) **(Fig 1D)**. We also quantified those residues that did not conform to any particular secondary structure throughout our simulation (existed as a secondary structure for < 70% of times throughout simulation). The percentage of these dynamical residues that make up each of the functional domains of the protein are shown in **Fig 1E**. These residues, displaying fast dynamics for inter-change of secondary structures, were found to predominantly exist in NBDs highlighting the underlying conformational flexibility of this region.

Additionally, it was observed that most of the helices that made up the protein were rigid (blue, helix angle close to 0) while a few of them displayed a fair degree of bending (red) **(Fig 1F)**. Incidentally, most of the helices showing flexibility lay within the TMDs. Helix flexibility has been shown to play a role in mechanisms of gating in ion channels(18). Since ABCA1 obtains its substrate laterally from the membrane, helical flexibility of this region may enable the substrate uptake mechanism.

### Volumetric changes in the tunnel enclosed within ABCA1

In order to characterize the tunnel enclosed within the protein, we generated a comparative volume as well as diameter profile of the Cryo-EM structure with respect to the simulation stabilized structure. The simulated structure appeared more extended and the differences in both the diameter as well as volume of the tunnel show drastic variations between the two structures in regions bounded by the TMDs and NBDs **(Fig 2 and S6)**. While the diameter of the pore enclosed within ECDs remains consistent in the two structures, there is significant divergence at two points (around 122 and 142 Å respectively, within the TMD region **(Fig 2)**.

**Figure 2:**
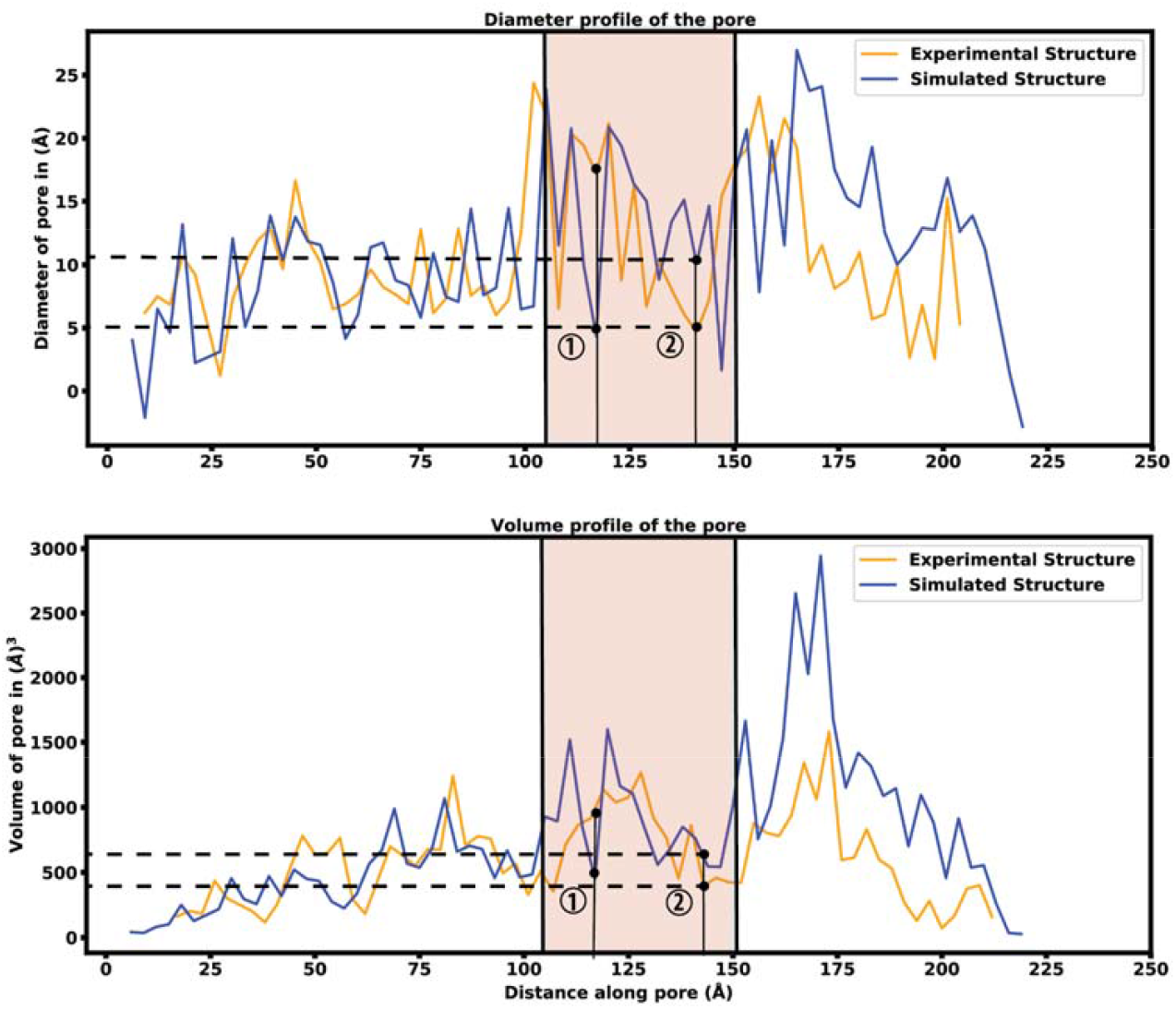
Characterization of the pore enclosed within ABCA1: A comparative diameter (top) as well as volume (bottom) profile showcasing the drastic variations observed within the Cryo-EM structure (orange) and simulation stabilized structure (blue).

The Cryo-EM structure reported a non-continuous central cavity in the TMD region of ABCA1(6) (**Fig 3A**). The two TMDs were found to contact each other through a narrow interface made by helix 5 and helix 11 of TMD1 and TMD2 respectively. This region at ∼142 Å in the volume profile was found to completely occlude the pore at the intracellular end **(Fig2 and 3A)**. This conformation of the protein therefore, resembled the outward-facing conformation of ABC transporters where an opening is present towards the exofacial side. Simulation stabilized structure however, shows a continuous tunnel that traverses the entire TMD region connecting the intracellular and extracellular edges **(Fig 2 and 3B)**. This central cavity in the TMD region of the simulated structure is wider at both its edges but forms a narrow aperture where helix 5 and helix 11 of TMD1 and TMD2 respectively, come close **(Fig S7A)**. This region, although hypothesized to form a substrate binding pocket was found to enclose a volume of 450 Å^3^ in the Cryo-EM structure which is too small to accommodate a single lipid (43). The simulation stabilized structure however, shows a significant widening in this region leading to an increase in its volume by about 1.5 times (∼654 Å^3^). Interestingly, since this region is bound by two flexible helices**(Fig 1F)**, changes in their orientation during the substrate export cycle may further alter the volume of this pocket This may also account for the versatility in substrate binding associated with ABCA1(19).

**Figure 3:**
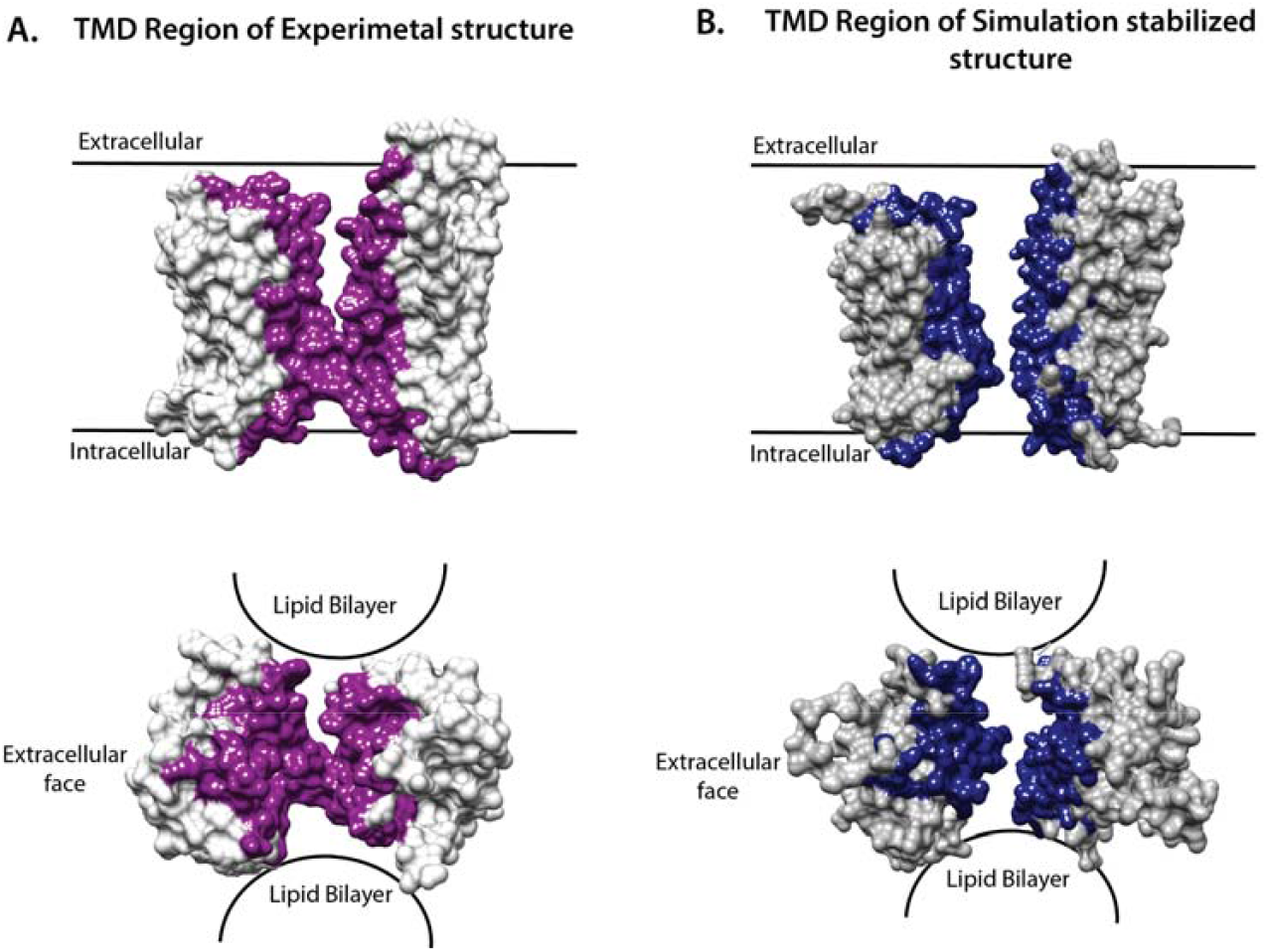
Tunnel enclosed within the TMD of ABCA1. Lipid exposed TMD1 and TMD2 of **A**. Cryo-EM structure and B. Simulated structure are depicted in surface representation highlighting the presence of a tunnel in this region. The top two panels represent the lateral view of the TMDs while the bottom two panels represent the TMD’s as viewed from the top. The experimental structure (top left and bottom left) shows a narrow opening while the simulation stabilized structure (top right and bottom right) shows a continuous tunnel from the intracellular end to the extracellular end. The surface colored in purple and blue represent residues that line the inner cavity.

Additionally, the region around 122 Å shows a dip in diameter in the simulation stabilized structure but not in the Cryo-EM structure. This region was formed by helix 5 and helix 7 of TMD1 and TMD2 respectively and has resulted due to a change in the orientation and tilt of these helices in the simulation stabilized structure. Although this region, too, is associated with a decrease in volume of the pore **(Fig 2)**, we believe that it would not hamper substrate transfer as it was found to be located towards the posterior end of the tunnel **(Fig S7B)**.

### Influence of ABCA1 on membrane curvature

In order to understand the fate of lipids in the heterogeneous membrane due to presence of ABCA1, we simulated another heterogeneous membrane with the same composition (see Materials and Methods for details) to serve as our experimental control. The average membrane thickness (of outer as well as inner leaflet) for both the membranes was computed. Our results indicated the membrane thickness to be fairly constant throughout all the five simulations as well as for control membrane **(Fig S8)**. The average thickness of the bilayer ranged from 41.2 - 41.9 nm for the membrane which had ABCA1 embedded in it, while that of the control membrane ranged from 41.7-42.6 nm.

**Fig 4A** depicts the time averaged deformation map for the top (average thickness ranged from -3.19 to 2.43 nm) as well as the bottom leaflet (average thickness ranged from -2.73 to 3.74 nm). This deformation map revealed that the outer leaflet experienced a positive deformation at the centre (region in red having thickness 1.24 to 2.43 nm) while the lower leaflet underwent a negative deformation (region in blue -0.34 to -2.73 nm) compared to the control membrane **(Fig S9)**. These striking differences in membrane thickness imply that the presence of ABCA1 leads the outer leaflet to be expanded while the lower leaflet condenses at the site of implantation of protein in the bilayer. Next, to ascertain whether any distortion was present in the membrane, mean curvature of both the leaflets along the membrane normal were calculated using a modified GridMAT algorithm. We observed that both the leaflets were found to be curved towards the middle of the bilayer at the points where the TMDs intersected with the bilayer **(Fig 4B)**. This indicates that the presence of protein indeed, induces a curvature in the bilayer. Interestingly, ABCA1 activity is believed to result in condensation of outer leaflet and expansion of inner leaflet which is in complete contrast with our observation of its resting state conformation(20). It is likely that this compression of the bilayer is necessary for lipid translocation(20).

**Figure 4:**
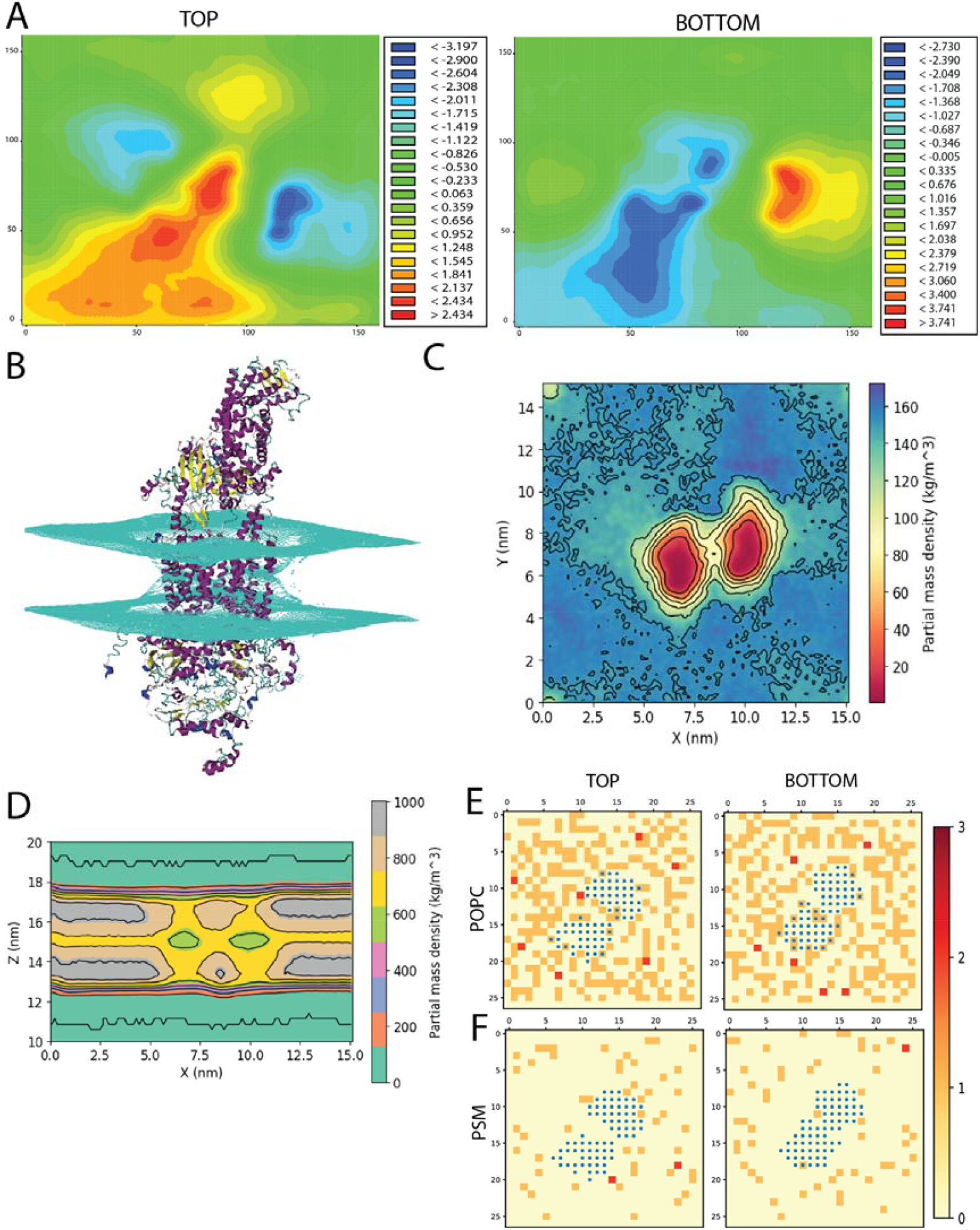
Effect of ABCA1 on membrane curvature and dynamics. **A**. Heat maps depicting average membrane thickness of the top and bottom leaflet of the membrane throughout simulation respectively. **B**. Membrane curvature induced in the membrane due to presence of protein. Average density of lipids across two leaflets along **C**. Z-axis and **D**. Y-axis. Frequency of lipid (shown as red, orange and cream) of **E**. POPC and **F**. PSM around protein (blue).

The average density profile of the simulation along Z-axis showed an overall reduction in lipids (normalized across the two leaflets) around the protein **(Fig 4C)**. The contours in the density profile computed along the Y-axis too showed that the membrane curves at the centre where the TMDs of ABCA1 are rooted **(Fig 4D and S10)**. Interestingly, there exists an island of densely populated lipids within the region between the two TMDs of the protein **(Fig 4E)**. This region may serve as the source of substrate for the protein.

### Influence of ABCA1 on lipid dynamics

Next, the average area spanned by the heterogeneous membrane with protein as well control membrane was computed as a measure of their stability throughout the simulation **(Fig S11)**. It can be observed that the average area spanned by the membrane with ABCA1 was a little higher than the control membrane (60.8 - 61.9 Å^2^ and 55.0 – 57.5 Å^2^ respectively).

In order to further explicate the protein-lipid interactions at play we performed clustering of each lipid type to generate separate heat maps based on lipid counts around the protein. **Fig S12 and 13** depicts the spatial location of each of the lipid types around the protein. It was observed that the most abundant lipid around the protein residues in both the top and bottom leaflet was POPC followed by PSM. **Fig 4E and F** illustrate the heatmap of PSM and POPC respectively where the gradient colours represent the number of lipid (PSM or POPC) molecules in the cluster and the blue squares represent the protein residues. While both the lipids did not appear to cluster as such, both POPC and PSM appeared to form a shell around the protein residues that were immersed in the membrane. We also determined the different lipid types that surrounded the entire protein within a radius of 0.5 nm throughout the simulation **(Fig 5B)**. We assumed that these lipid residues may be interacting with the corresponding protein residues owing to their close vicinity. Our results indicated that the most prevalent lipid type within the shell radius of the protein is POPC followed by PSM.

**Figure 5:**
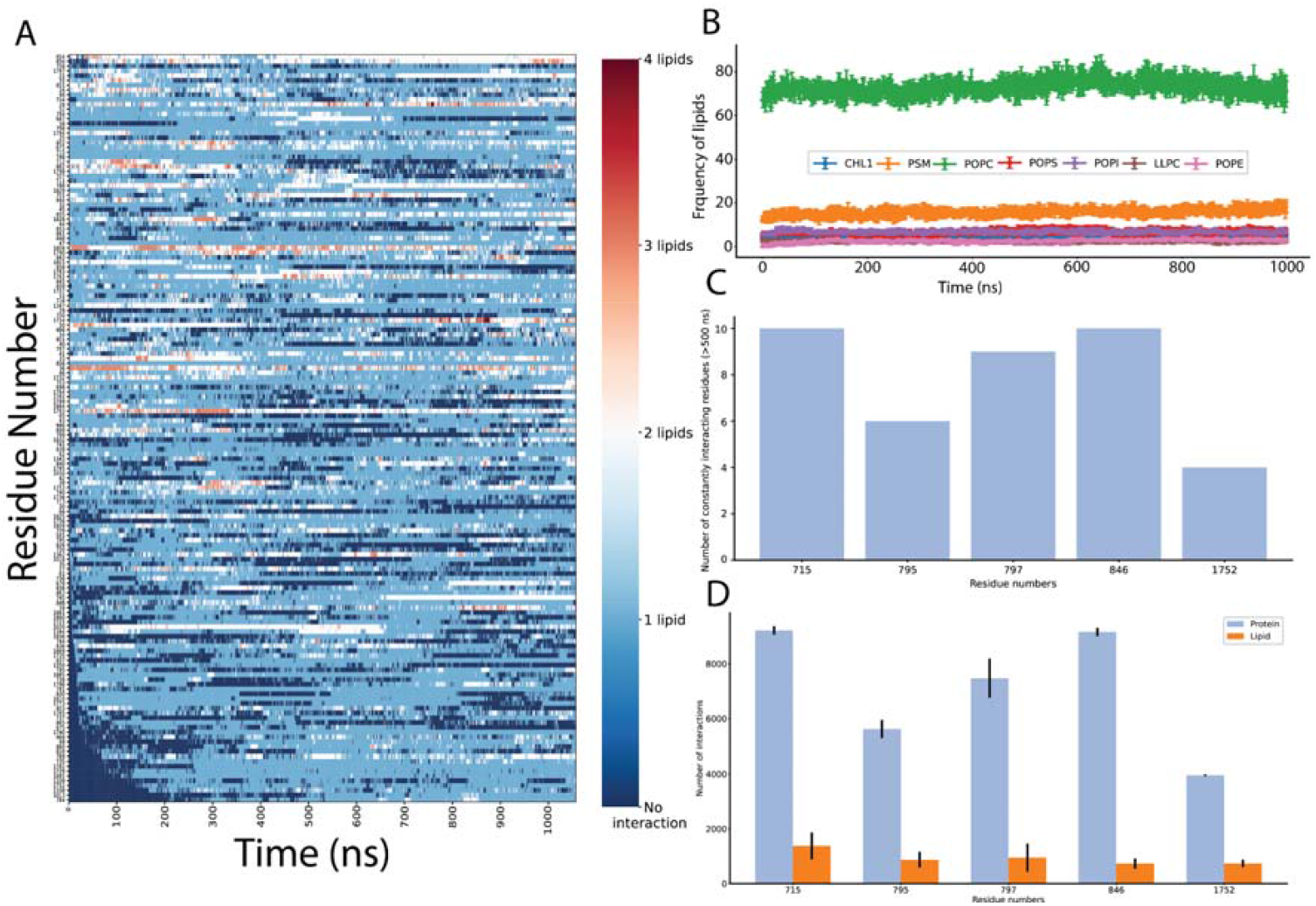
Protein-lipid interactions’ analysis. **A**. Frequency of each lipid type surrounding the protein residues within a radius of 0.5nm throughout our simulation **B**. Interactions for each of the protein residue with lipids (0 or more) throughout the simulation **C**. Number of protein-protein contacts made by the five disease-related protein residues **D**. Frequency of protein-protein and protein-lipid interactions of the five disease-related protein residues

### Elucidating the protein-lipid interactions that stabilize ABCA1 in the membrane

In order to study the protein-lipid interactions, we focussed on the protein residues that were found to interact with these surrounding membrane lipids for more than 500 ns throughout the simulation. A total of 156 such residues were found and the total number of interactions for each of these residues was quantified **(Fig 5A)**. It was observed that some of these residues did not interact with any lipid residues initially (depicted in dark blue at the bottom of the heatmap), but were found to form transient interactions with 1-4 lipids as the course of simulation progressed. All such protein-lipid interactions were aggregated and have been depicted according to lipid types that they interact with in **Fig S14**.

Subsequently, we analysed, if any of the 156 residues that were found to actively interact with lipids throughout our simulation were implicated in any of the ABCA1 related disorders to extrapolate the importance of their interactions with surrounding lipids in protein stability or function. Those residues amongst these 156 that were reported to be pathogenic or disease-related in either UniProt or dbSNP or both were assessed. Furthermore, SNPs & Go(21) were also used to ascertain the disease-association of the shortlisted nsSNPs. SNPs & Go results include, in addition to itself, predictions from PhD-SNP(22) and PANTHER(23). 5 amongst the 156 residues (L715del, T795P, K797N, V846I, S1752C) were identified to have high probability to be disease causing. These were associated with pathogenic phenotypes such as increased cardiovascular risk & increased plasma cholesterol levels. **Fig 5C** depicts the intensity of protein-protein contacts of these 5 residues while **Fig 5D** enumerates the number of protein-protein as well as protein-lipid interactions that these 5 residues participated in. Additionally, it was also observed that these residues clustered mostly in TMD1 where it made contact with the outer as well as inner leaflet. They were also found to interact with POPC and PSM mostly **(Fig S14)**. Our results indicate that these residues are involved in several stabilizing protein-protein as well as protein-lipid interactions and therefore, mutations in these regions may potentially be destabilizing in nature and/or affect protein function.

### Conclusion

The results presented herein represent the first simulation of ATP unbound monomeric ABCA1 in its native heterogeneous membrane environment. The simulation stabilized structure so obtained deviates from the Cryo-EM structure, namely in the volume of the tunnel enclosed within its TMDs and NBDs. This structure also shows presence of a continuous central cavity that traverses the entire protein which was missed in the Cryo-EM structure. The presence of cavity in TMD region forms a narrow aperture where helix5 and helix 11 of TMD1 and TMD2 respectively come close. Helix 5 and helix 11 show a high degree of flexibility and their orientation is suggestive of this site serving as a lipid binding pocket. Moreover, this pocket may undergo changes in volume to allow binding of a variety of lipid substrates.

We have also documented the protein-lipid interactions that detail the effect of protein on membrane dynamics at an atomistic resolution. Taken together our observations suggest that the presence of ABCA1 exerts a differential role on leaflet curvature due to which both the leaflets curve inwards at the site of implantation of the protein. The lipids were also found to be sparsely packed around the protein (decreased overall density), when the protein was in its inactive state (ATP unbound).

Recently, reverse cholesterol transport has surfaced as a focal point for regulation of lipid metabolism, especially given its role in cardiovascular disease. Since ABCA1 mediated lipid efflux is the first step of the pathway, establishing its stable conformation and mechanism of action is of immense importance. In summary, our study has refined the original model of the ATP unbound ABCA1 by studying it in its native environment and these novel findings will pave the way for future studies.

## Computational Methods

### Structure preparation and Classical Molecular Dynamics

CryoEM structure of ABCA1 with an overall resolution of 4.10 A (PDB ID: 5xjy) was modelled for missing regions using Modeller(24) and inserted into a heterogenous membrane. The heterogeneous membrane was made of 1-palmitoyl-2-oleoyl-sn-glycero-3-phosphocholine (POPC) (482 molecules), palmitoyl sphingomyelin (PSM) (108 molecules), Cholesterol (CHL) (48 molecules), 1-palmitoyl-2-oleoyl-sn-glycero-3-phosphoinositol (POPI) (36 molecules), 1-Palmitoyl-2-oleoyl-sn-glycero-3-phosphoethanolamine (POPE) (30 molecules) and 1,2-palmitoyl-oleoyl-sn-glycero-3-phosphoserine (POPS) (24 molecules), Lysophosphatidylcholine (LLPC) (18 molecules).

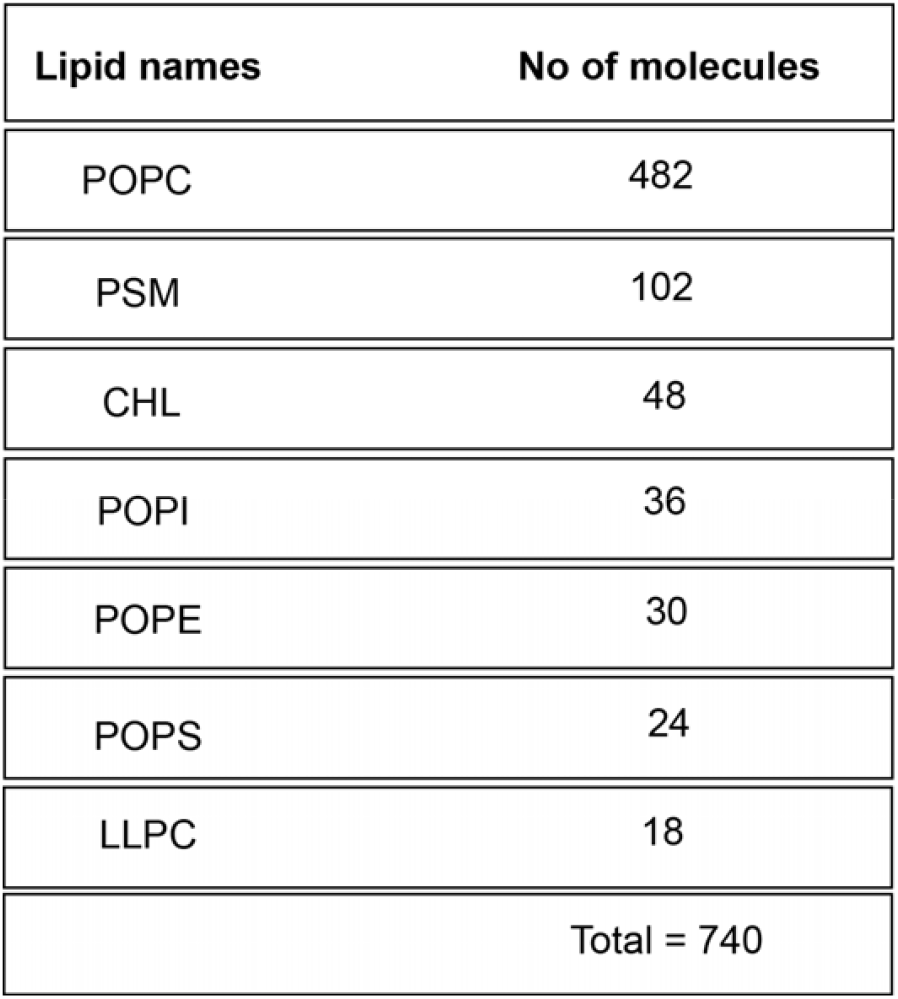

MD simulations were performed with the GROMACS 4.6.1(25–27). The initial system was built using CHARMM-GUI and is represented by CHARMM36 all-atom force-field(28–30). The water was modeled using the TIP3P representation(31). Each of the five starting conformations were placed in a dodecahedral water box (volume = 6477.957 nm^3^) large enough to contain the protein-membrane complex and at least 1.0 nm of solvent on all sides. Periodic boundary conditions were used, and the long-range electrostatic interactions were treated with the particle mesh Ewald method(32) using a grid spacing of 0.12 nm combined with a fourth-order B-spline interpolation to compute the potential and forces in-between grid points. The real space cutoff distance was set to 1.2 nm and the Van der Waals cutoff to 1.2 nm. The bond lengths were fixed(33), and a time step of 2 fs for numerical integration of the equations of motion was used. Coordinates were saved every 200 ps. The pressure coupling was done by employing a Parrinello-Rahman barostat(34) using a 1 bar reference pressure and a time constant of 5.0 ps with compressibility of 4.5e-5 bar using semi-isotropic scaling scheme. Eighty-two positive counter-ions (K+) were added, replacing eighty-two water molecules so as to produce a neutral simulation box. All the starting structures were subjected to a minimization protocol for 5000 steps using the steepest descent method followed by six sequential equilibration steps with restraints on the protein atoms as per the CHARMM-GUI protocol.

Five independent trajectories, each of 1 μs at 300 K were carried out. The combined time-scale of our simulations is 5 μs.

A control membrane (without the protein) with same composition (POPC: 482, PSM: 102, CHL: 48, POPI: 36, POPE: 30, POPS: 24, LLPC: 18) was similarly constructed using CHARMM-GUI using the CHARMM36 force field. One hundred and forty-one positive counter-ions (K+) as well as eighty-one negative counter-ions (Cl-) were added, replacing two hundred and two water molecules to produce a neutral simulation system. This system was subjected to minimization and equilibration as previously mentioned, in accordance with CHARMM-GUI protocol. The control membrane was similarly placed in a water box large enough to contain it with at least 1.0nm of solvent on all sides. Periodic boundary conditions were used, and the long-range electrostatic interactions were similarly treated with the particle mesh Ewald method using a grid spacing of 0.12nm combined with a fourth-order B-spline interpolation to compute the potential and forces in-between grid points. The real space cutoff distance was set to 1.2 nm and the Van der Waals cutoff to 1.2nm. The bond lengths were fixed, and a time step of 2 fs for numerical integration of the equations of motion was used. Coordinates were saved every 500 ps. The pressure coupling was done by employing a Parrinello-Rahman barostat using 1 bar reference pressure and a time constant of 5.0 ps with compressibility 4.5e-5 bar using semi-isotropic scaling scheme.

### Simulation Analysis

#### Domain assignment

The six domains in ABCA1 structure were assigned and colored accordingly using UCSF Chimera 1.14(35). The domains were assigned in accordance with the structure of the human lipid exporter ABCA1(6). The range of residues and the assigned color(s) are as follows: ECD1 (46 to 630, cyan), ECD2 (1366 to 1640, orange), TMD1 (1 to 45 and 631 to 902, blue), TMD2 (1327 to 1365 and 1641 to 1906, lime), NBD1 (903 to 1147, yellow), and NBD2 (1907 to 2143, magenta).

#### Domain motion analysis

Principal component analysis was performed using the covariance matrix of the atomic coordinates throughout the simulation. Eigenvectors (representing modes of fluctuation) and eigenvalues were obtained by diagonalizing this matrix. Eigenvector 1 representing the most dominant mode of motion was used to produce a PDB file containing 100 conformations along this vector. This was visualized in the Normal Mode Wizard of VMD(36).

#### Free energy estimation and clustering

InfleCS was used to estimate the density and free energy using Gaussian mixture models (GMM)(37). A 2D array of RMSD and Rg of all five sims was given as the input. The estimated density was then clustered using InfleCS clustering and the free energy was visualised. The core states were identified at density maxima using the estimated Gaussian mixture density and the points were divided into clusters.

#### Helix flexibility

VMD Bendix plugin(38) was used to calculate and visualize both dynamic and static helix geometry. RWB or Red-White-Blue heatmap colors are used for visualisation. Red indicates the areas of highest helix axis angle (>15□), followed by white for intermediate angle magnitudes (0-15□) and blue for angles close to, or at 0□. Therefore, red-colored regions are highly flexible while blue regions are rigid. The Bendices ends that are at angle of 0 per definition are not resolved by the axis-generating algorithm.

### Characterization of the Inner tunnel enclosed within ABCA1

The crystal structure (PDB ID: 5xjy) as well as the simulation-stabilized structure were submitted to the CICLOP webserver(39) for a complete characterization of the inner residues lining the pore. The structures were first aligned using Chimera(35) and then submitted under manual mode of alignment and the diameter and volume profiles generated by the tool were used as is. CICLOP detects and reports the protein atoms that surround the inner cavities.

#### Average lipid thickness

Lipid thickness across both leaflets was calculated after equilibration using the MEMBPLUGIN(40) software. MEMBPLUGIN was used for calculating average membrane properties such as the membrane thickness. The Membrane Thickness tool contained in this software calculates the average membrane thickness over a chosen trajectory by measuring the distance between the two density peaks of user-selected atoms (phosphorus) belonging to the head group of phospholipids. The average lipid thickness was calculated for all five sims post stabilization. The mean average thickness across all five simulations was then plotted.

#### Membrane partial density

This metric was calculated using g_mydensity on a grid with spacing of about 0.2 nm(41). Therefore, each element of volume (voxel) was ∼ 0.2 nm × 0.2 nm × *Z* nm, where *Z* is the dimension of the box in the direction of the membrane normal. Partial densities were calculated as the average mass of the particles present in the voxel divided by the volume of the voxel; only lipids were taken into account.

#### Thickness maps

Lipid thickness maps for both leaflets were generated via MEMBPLUGIN(40) software. Time-averaged deformation maps were generated by extrapolating the z coordinates of P atoms into a regular orthogonal grid in the XY plane. Irrespective of the leaflet, deformation is positive when it expands the membrane, and negative otherwise. The thickness maps were visualized in a contour plot.

#### Lipid clustering

Cluster analysis was done using an in-house Python script. The last frame of the simulated trajectory was extracted in the PDB format. The PDB file thus generated was parsed to extract all the relevant information for all the atoms in the system. For each lipid/protein residue, a mean position was calculated by taking the arithmetic mean of the x, y, and z coordinates of the atoms pertaining to the lipid/protein residue in question. Taking the mean z coordinate of the thus generated coordinates, the membrane bilayer was split into two monolayers i.e. the top and the bottom layer. The desired layer was then further divided into square sections of 6 Å length. For each such box, the number of residues of the desired lipid class lying inside the box were calculated and plotted. At the same time, the protein residues lying inside the desired monolayer were calculated using the z coordinates as an indicator of the same. For all the protein residues thus identified, a similar estimation of a single coordinate was done similar to the way it was done for lipid residues. Finally, all the little 6 Å boxes containing at least 1 such protein residue were marked on the plot with a blue square while the lipid residues were marked with colors according to their prevalence (frequency).

#### Bilayer curvature

Further, to check whether the lipid membrane was curved to any extent because of the presence of the protein and to calculate the mean curvature of a membrane leaflet, the g_lomepro suite(42) was used. To determine the mean curvature, the lipid coordinates along the bilayer normal are first mapped onto a grid space, using a modified version of GridMAT-MD algorithm. Subsequently, a filter function was used to transform the grid-mapped coordinates to Fourier space and were then recovered via an inverse transform. First- and second-order derivatives of the filtered coordinates were used to calculate the mean curvature. In the current study, the ABCA1 protein was centred, and the last frame of each trajectory was analyzed. The mean curvature was individually analyzed for each set. The phosphate head groups were used to represent the membrane. A leaflet is defined to be positively curved if it bends away from the bilayer centre toward ABCA1 and negatively curved if it bends toward the bilayer centre away from ABCA1. Further, the visualisation of the curved structures was done using VMD(36) by keeping bin-x and bin-y parameters as 100.

## Author Contributions

**Sunidhi:** Methodology, Data analysis, Writing-Original draft preparation. **Sukriti Sacher:** Methodology, Data analysis, Writing-Original draft preparation, Reviewing and Editing. **Parth Garg:** Software. **Arjun Ray:** Conceptualization, Methodology, Supervision and Writing-Review and Editing.

## Acknowledgement

The authors would like to thank the HPC facility of IIIT Delhi for the computational facility.

