## Supplementary figures for "Elucidating the structural features of ABCA1 in its heterogeneous membrane environment"

### Elucidating the structural feature of ABCA1 in heterogeneous membrane environment - Supplementary Data

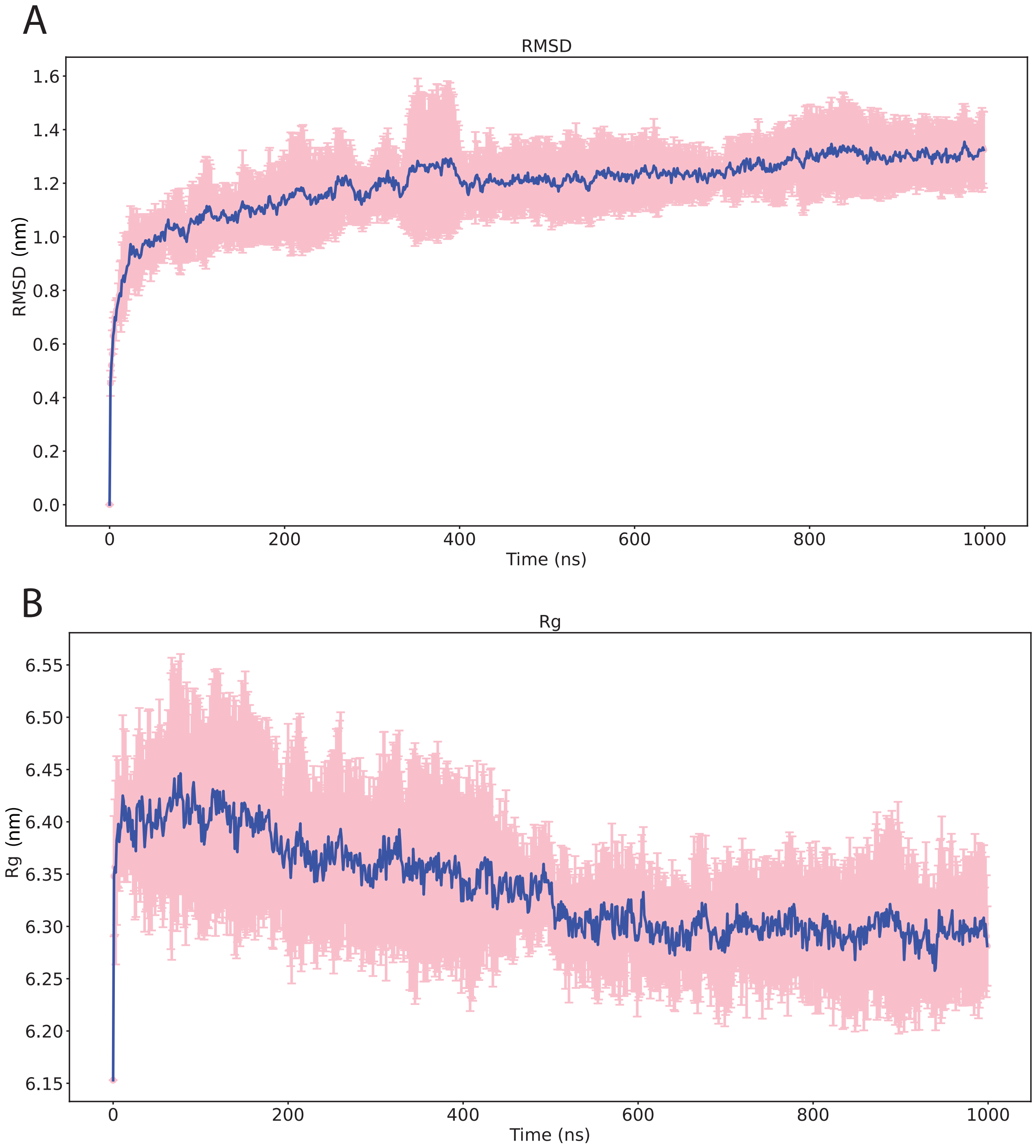


Figure 1: A. Root mean square deviation of backbone of protein across all five simulation. The blue line represents the average while error bars represent the mean absolute deviation observed along simulation time. B. Radius of gyration of protein across all simulation where green line represents the average and error bars depicts mean absolute deviation.

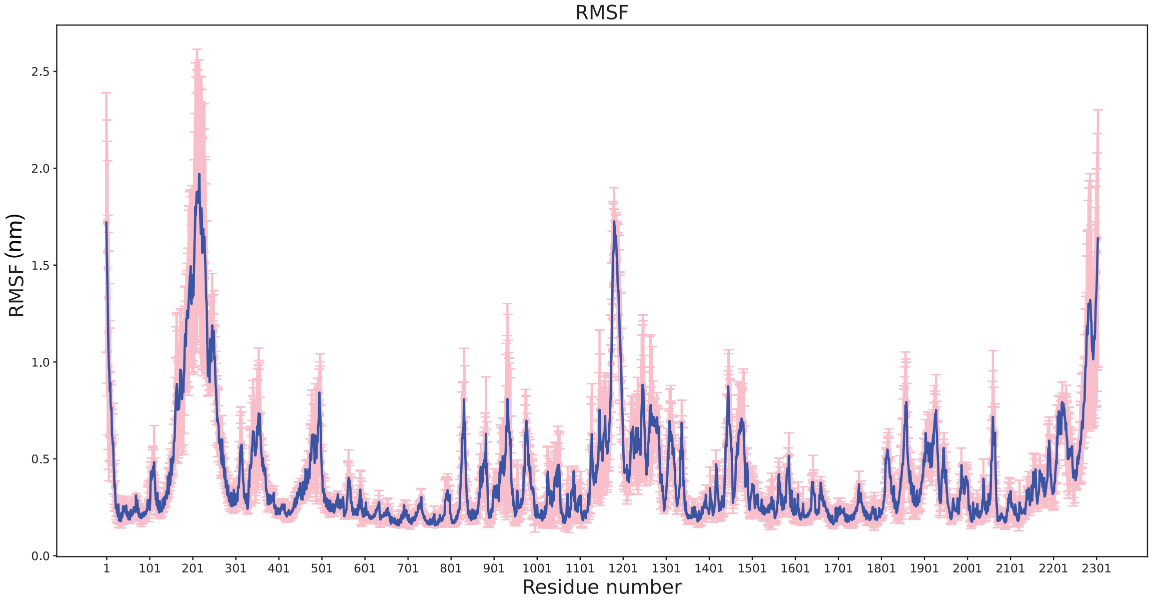


Figure 2: Residue-wise fluctuation (RMSF) observed in all the five simulations. The blue line represents the average while the error bars represent the mean absolute deviation.

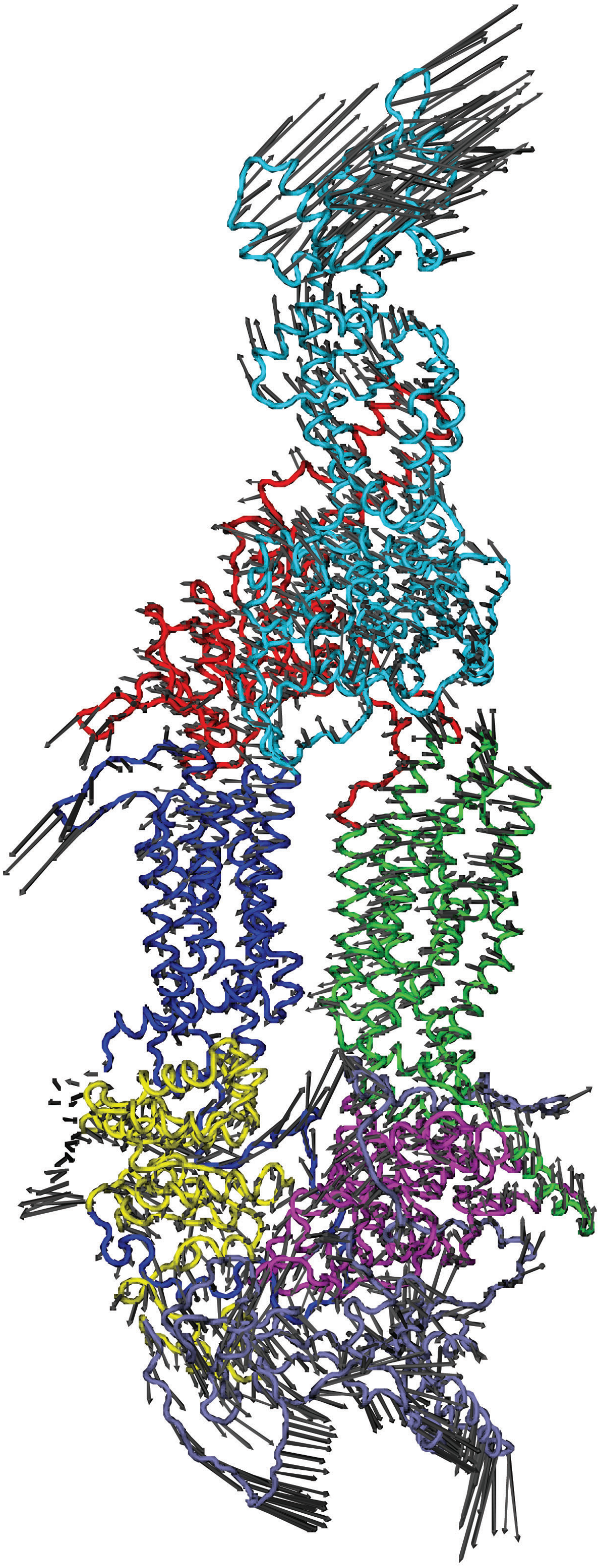

Figure 3: Domain concerted motion of ABCA1. Principal component analysis showing the dominant mode of motion of different domains of ABCA1 throughout the simulation. The direction of arrows (gray) depict the direction of motion of the domain, while their length depicts the intensity. The six domains that make up the protein include ECD1(cyan), ECD2(red), TMD1(blue), TMD2(green), NBD1(yellow), NBD2(magenta) and unspecified region located intracellulary shown in grey color.

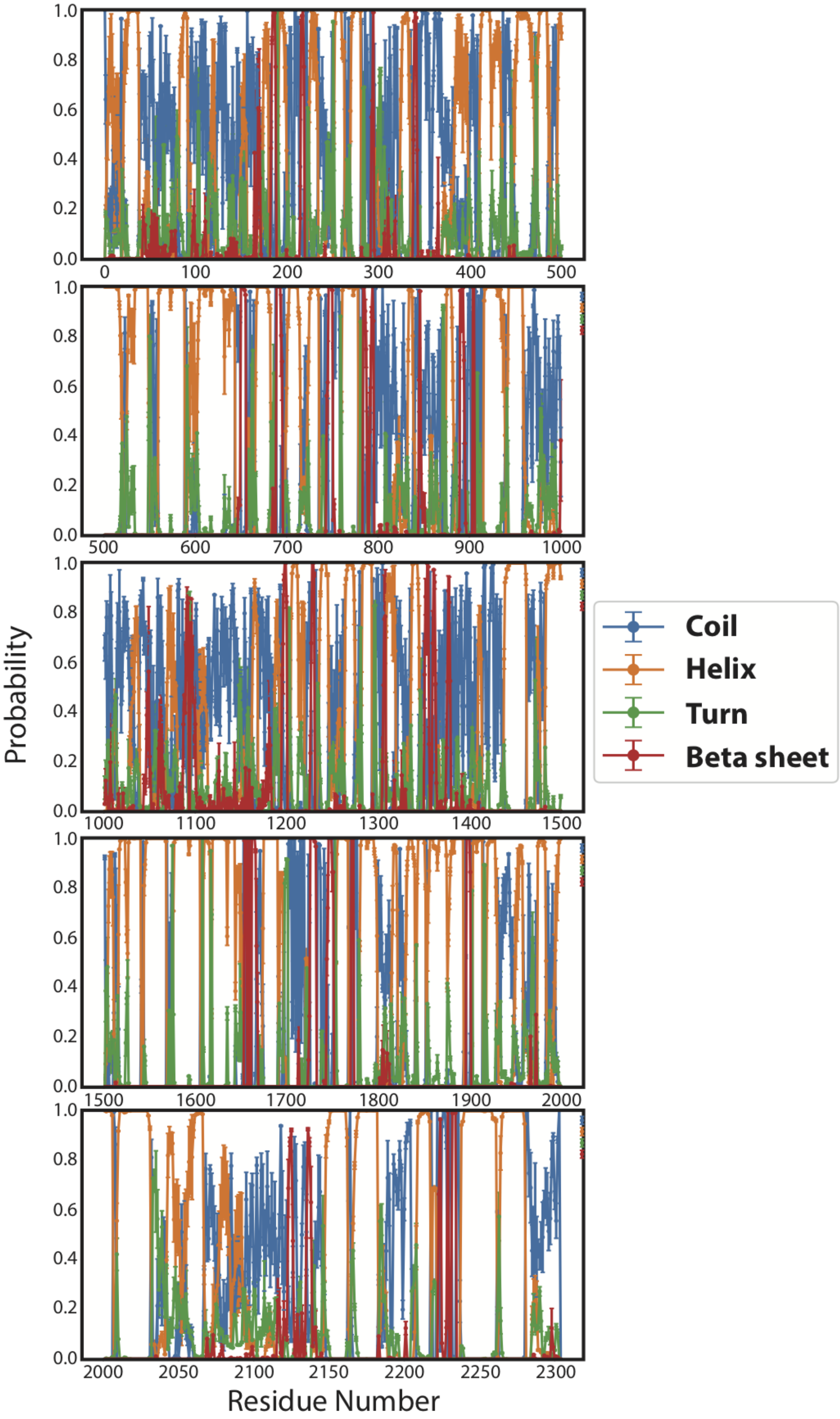


Figure 4: Propensity of each residue of the protein to conform to a particular secondary structure.

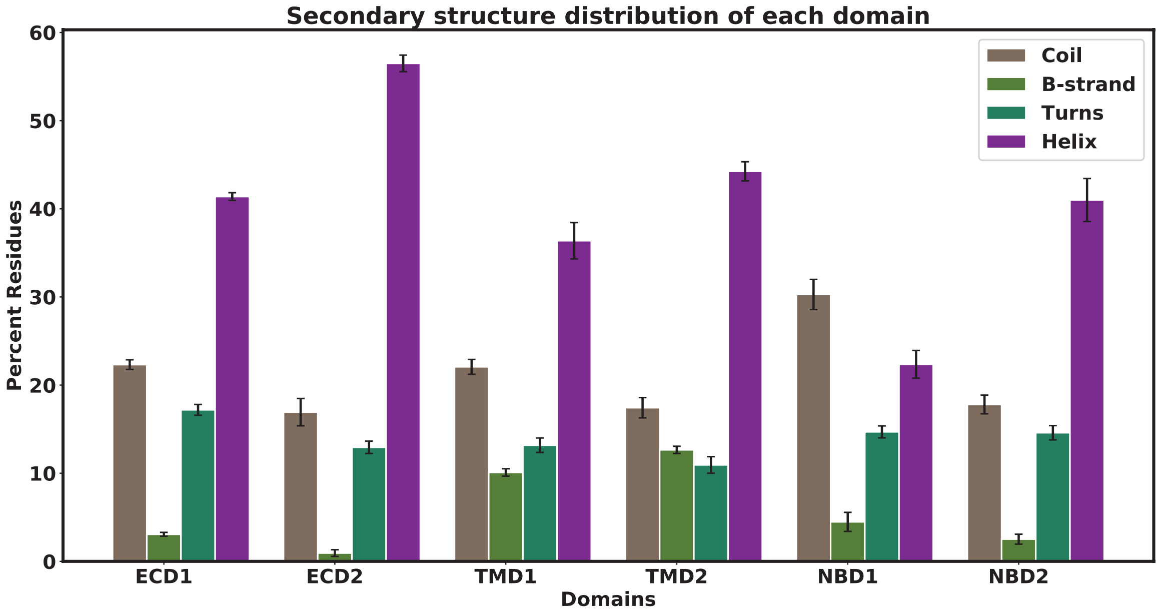


Figure 5: Secondary structure distribution for each domain of ABCA1.

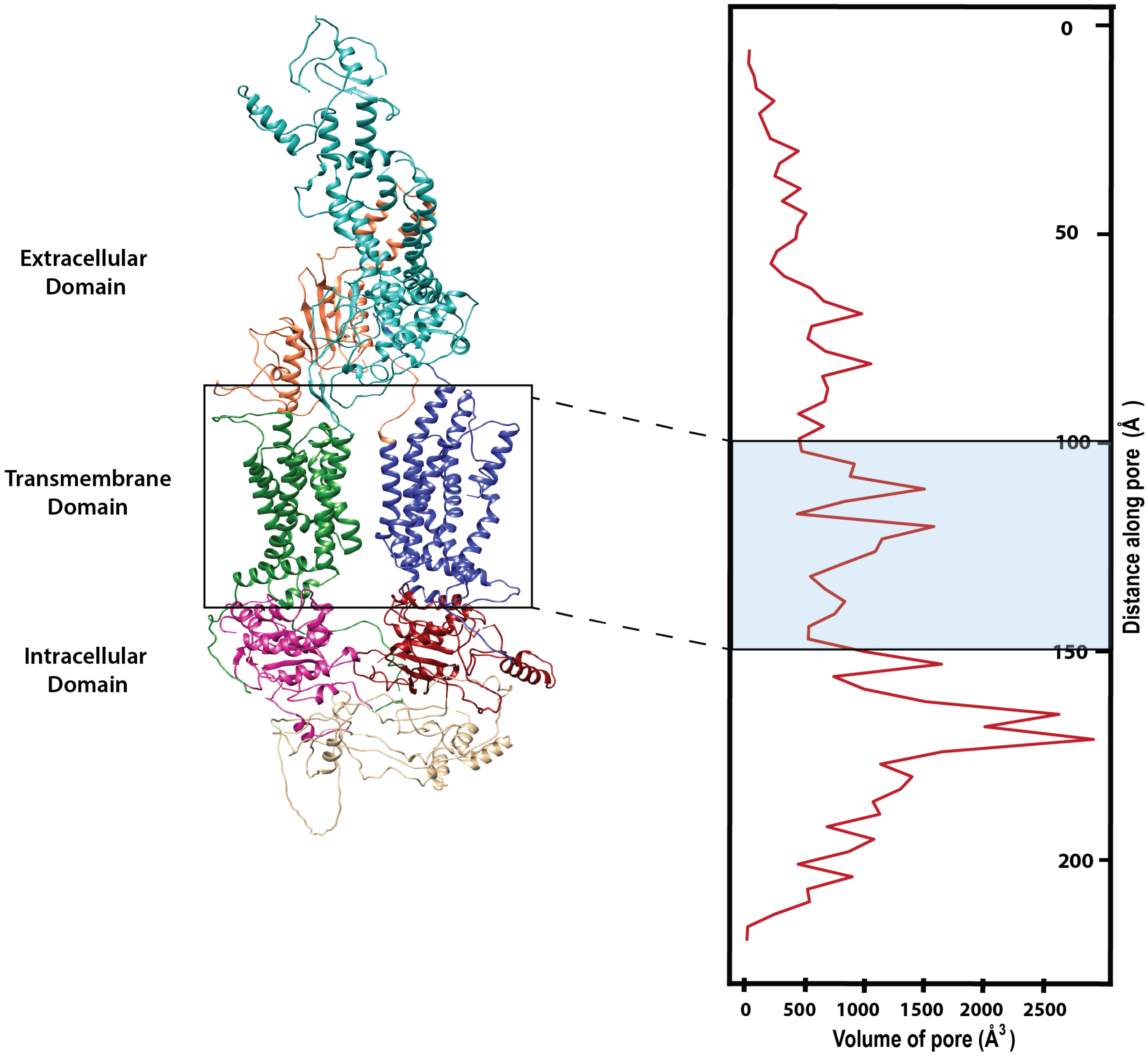


Figure 6: Characterization of the inner cavity lining of ABCA1. Hydrophobicity profile of the pore enclosed within simulation stabilized structure (left) with the simulation stabilized structure (center) showcasing the various domains : ECD1 (cyan), ECD2 (orange), TMD1 (green), TMD2 (blue), NBD1(magenta), NBD2 (brown). A comparative volume profile of the pore enclosed within simulation stabilized structure as well as the Cryo-EM structure has also been presented (right). The region on the profile that is buried within the membrane has been highlighted.

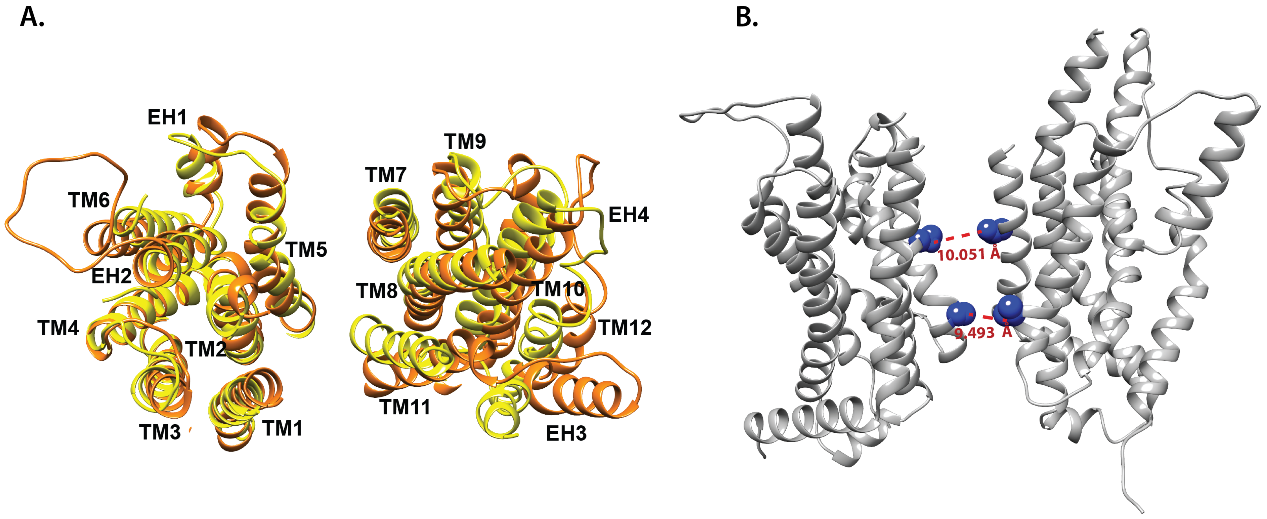


Figure 7: Transmembrane domain of ABCA1: A. TMD region of Cryo EM structure (yellow) superimposed onto TMD region of simulation stabilized structure (orange). The various helices TM1-6 and EH1-2 that constitute TMD1 and TM7-12 and EH3-4 that make up TMD2 are shown. B. Diameter of the pockets formed due to orientation of helices of TMD1 and TMD2. The points marked as 1 (10.051 Å) and 2 (9.493 Å) in the pore profiles shown in Figure 2 have been highlighted.

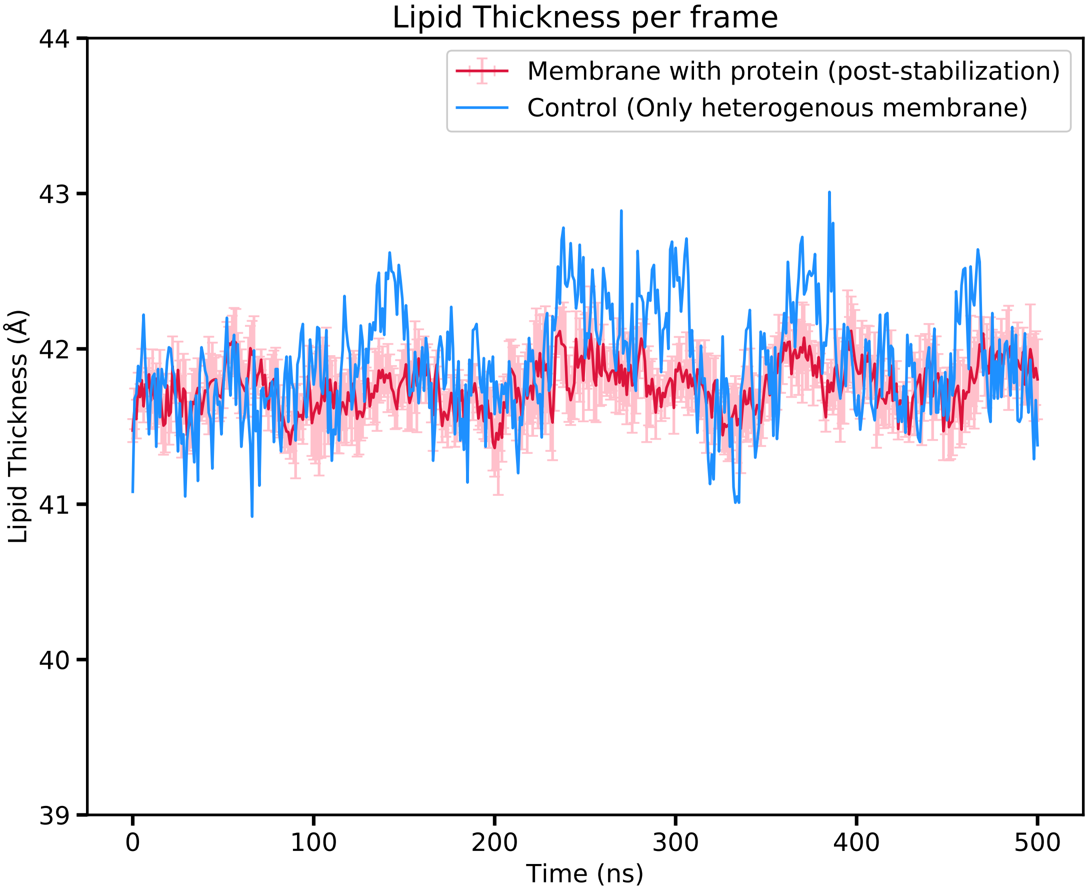


Figure 8: Membrane thickness across the simulation time. Thickness averaged across the top and bottom leaflet has been plotted for the control membrane (blue) as well as membrane with ABCA1 embedded in it (red). Error bars depict the mean absolute deviation in thickness across all five simulations.

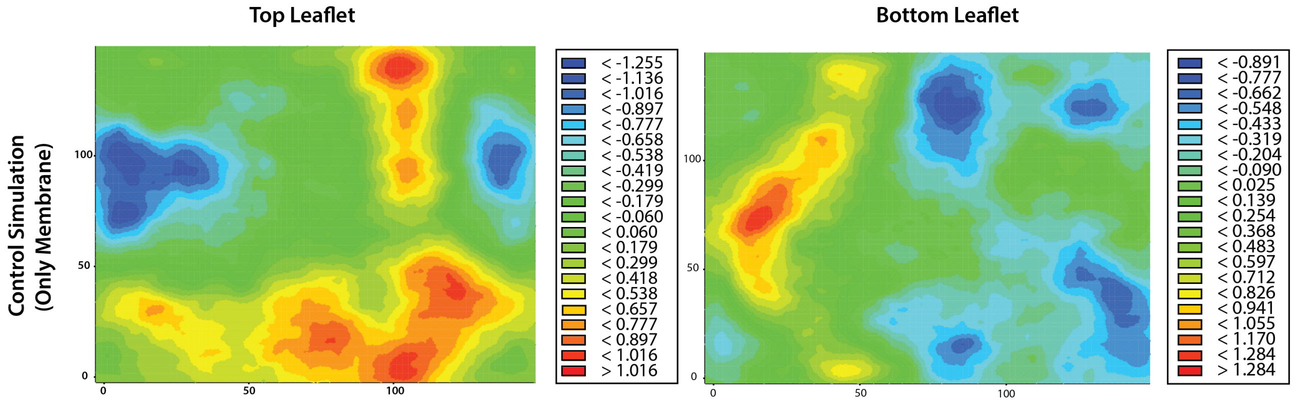


Figure 9: Average thickness map of the top leaflet (left) as well as bottom leaflet (right) for the control membrane.

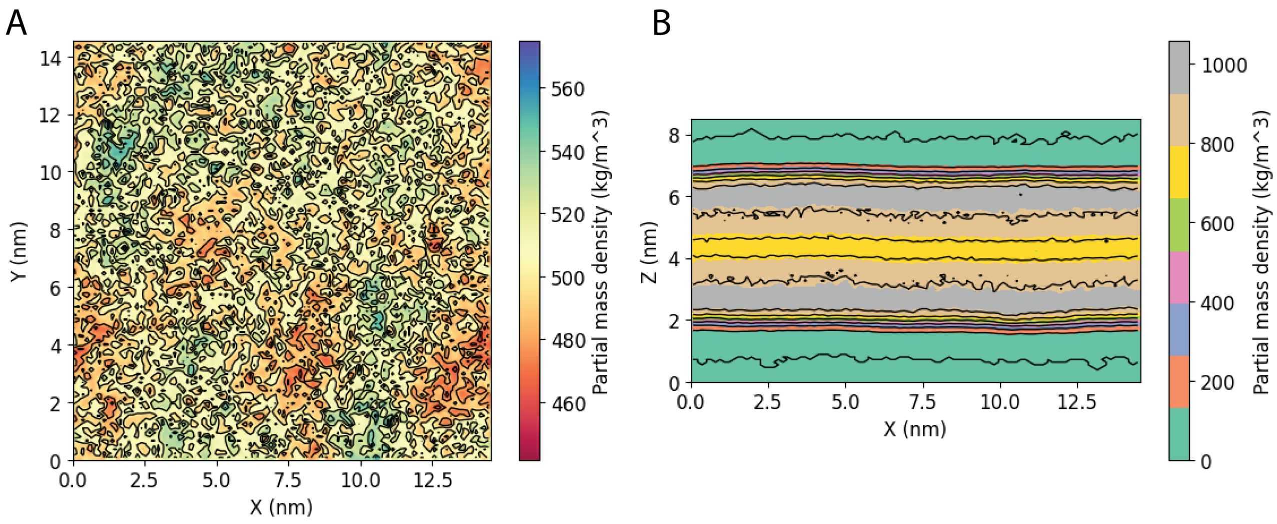


Figure 10: Average density profile of the control membrane computed along A. Z-axis and B. Y-axis.

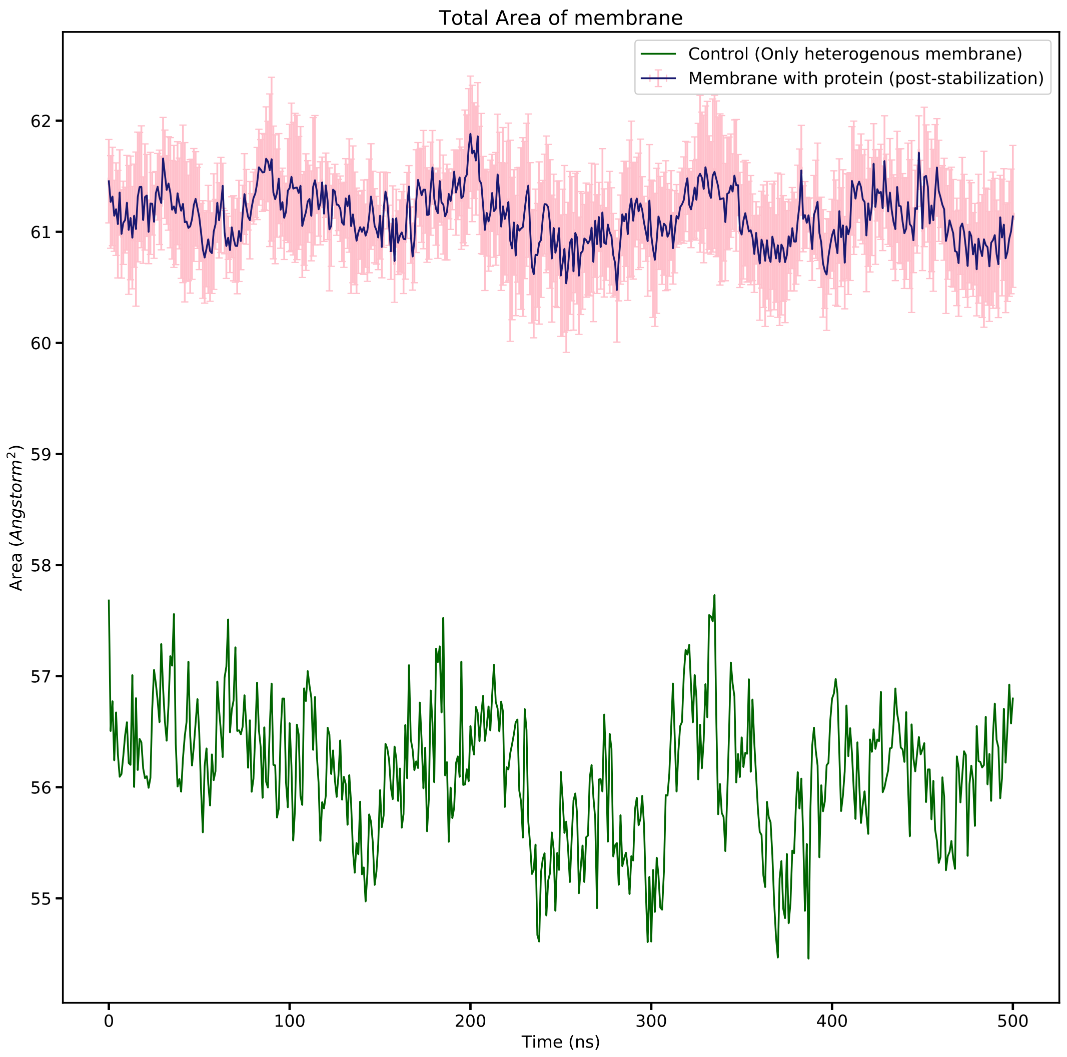


Figure 11: Total area averaged across the top and bottom leaflet for control membrane (green) as well as the membrane with ABCA1 embedded in it (blue). Error bars represent the mean absolute deviation in total area across all five simulations.

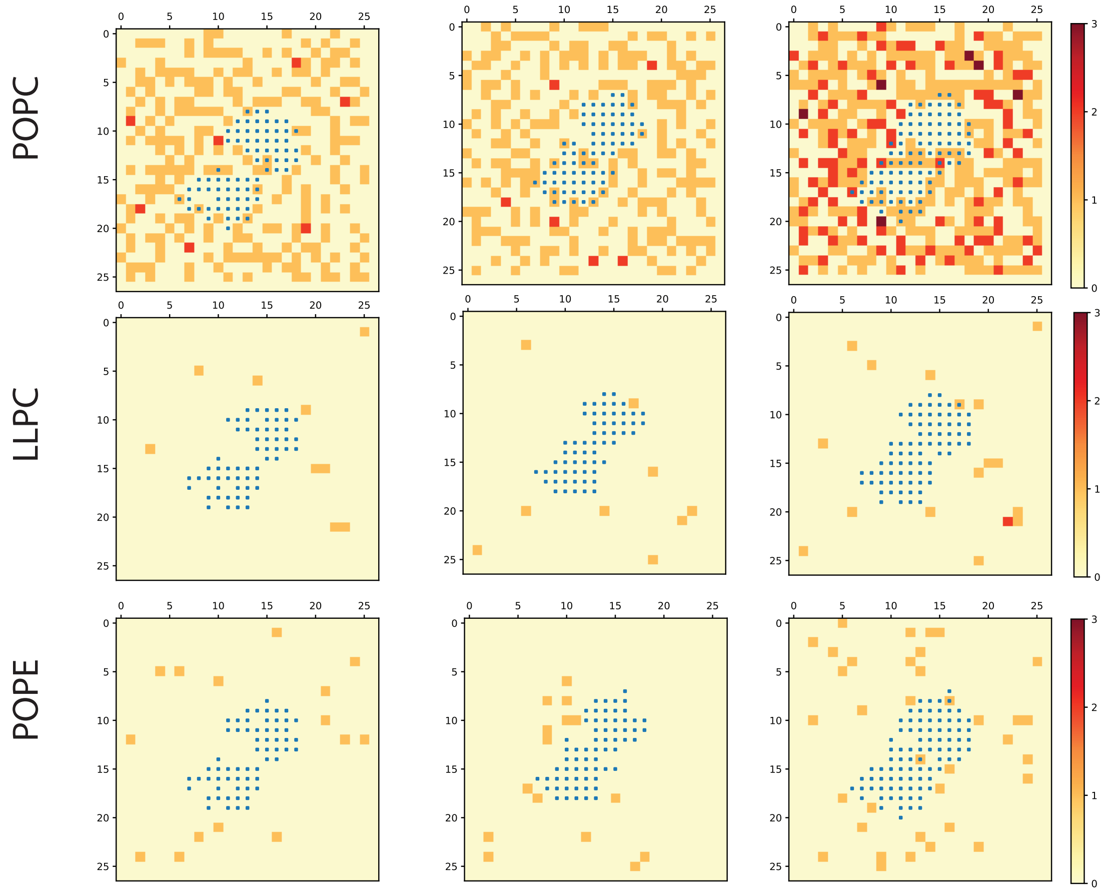


Figure 12: Lipid clustering. Lipids (POPC, LLPC, POPE) colored either as red, orange or cream depending on their frequency around protein (blue squares).The frequency for each lipid was computed for both the top and bottom layer separately and have been shown along with the combined map that represents the total frequency across both leaflets.

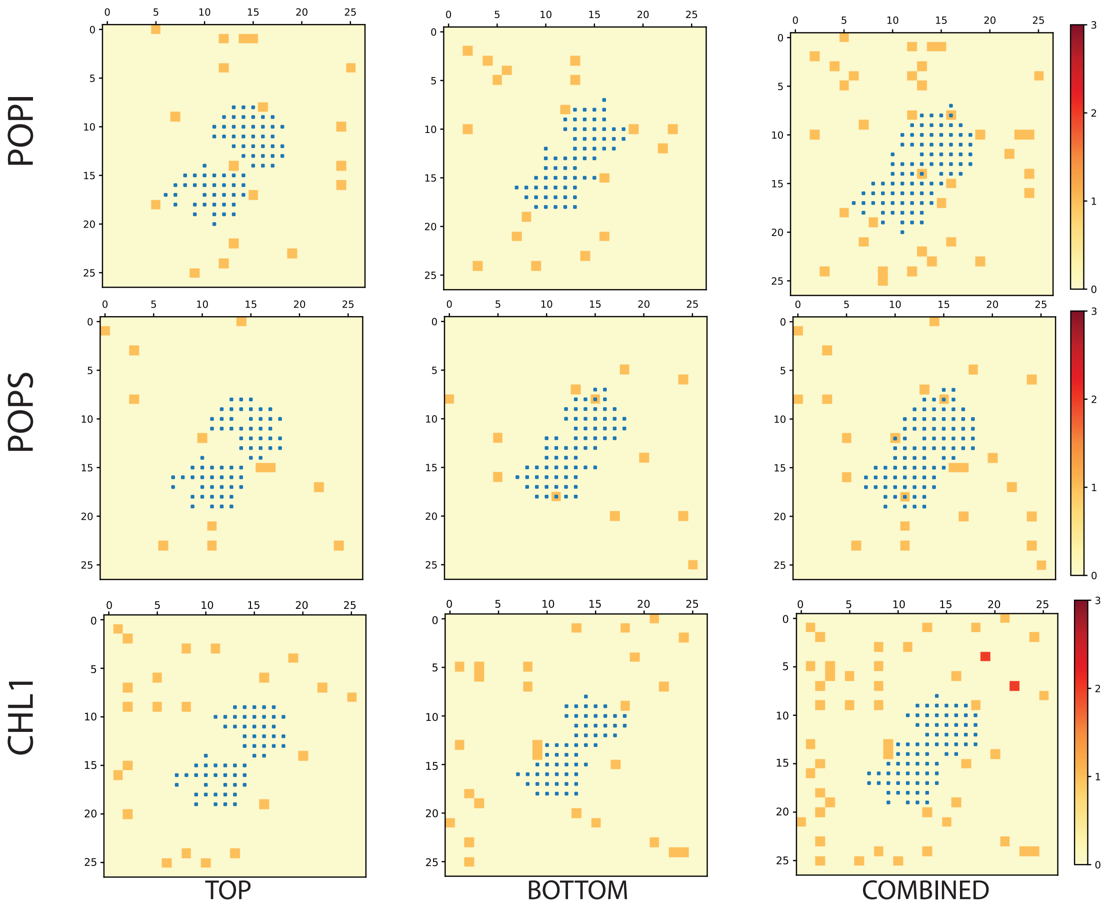


Figure 13: Lipid clustering. Lipids (POPI, POPS, CHL) colored either as red, orange or cream depending on their frequency around protein (blue squares). The frequency for each lipid was computed for both the top and bottom layer separately and have been shown along with the combined map that represents the total frequency across both leaflets.

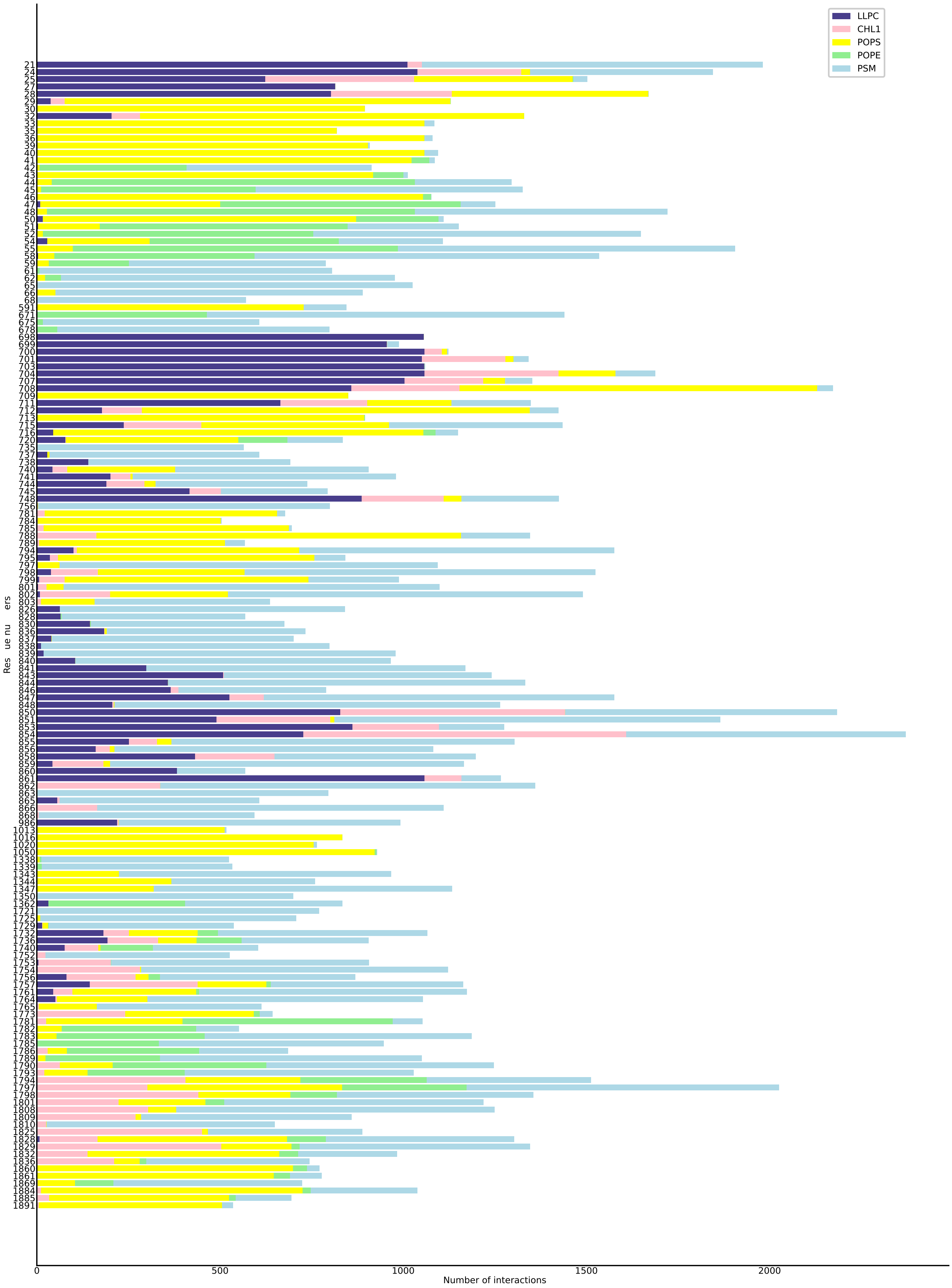


Figure 14: Number of interactions between protein residues in contact with lipids. Frequency of interaction for those protein residues that remained in contact with lipids for half the simulation time (500ns) has been depicted.
